# MR-Guided Focused Ultrasound Neuromodulation of Nucleus Accumbens to Attenuate Oral Fentanyl Motivation in A Self-Administration Rat Model

**DOI:** 10.64898/2026.10.17.759605

**Authors:** David A. Martin, Abubakr Eldirdiri, Thomas Ernst, Eric Cunningham, Segun Bernard, Sandesh Kamdi, Donna J. Calu, Linda Chang

## Abstract

**Introduction:** Transcranial low-intensity focused ultrasound (LIFU) is being explored as a non-invasive approach for neuromodulation to treat opioid use disorder (OUD).

**Methods:** We evaluated MR-guided focused ultrasound, with LIFU, targeting nucleus accumbens (MRgLIFU-NAc) bilaterally, using MR-acoustic radiation force-imaging (MR-ARFI) to ensure accurate targeting and adjust for FUS intensity delivered at the target. Male and female rats were trained to self-administer fentanyl/sucrose (70ug/ml fentanyl citrate in 5% sucrose) or 5% sucrose alone. Three cohorts of rats (total n=76) were evaluated on progressive ratio (PR) responding for oral fentanyl/sucrose at either 4 or 24 hours post-MRgLIFU-NAc. Animals in the first two cohorts received repeated rounds of LIFU or sham (anesthesia-duration matched). In cohort 3, we used a counterbalanced within-subjects design in which rats received both LIFU and sham one week apart.

**Results:** Reducing sucrose concentrations decreased responding for sucrose alone more than for fentanyl/sucrose, demonstrating a role for fentanyl in the reinforcing properties of fentanyl/sucrose. Across three sonication schemes, MRgLIFU-NAc did not attenuate PR performance for oral fentanyl/sucrose at 4 or 24-hours post-LIFU relative to controls. Adjusting LIFU intensity with MR-ARFI did not correlate with changes in PR from baseline. We confirmed that NAc is the correct target by bilateral inactivation of NAc with muscimol/baclofen mixture (50 + 50 ng/side), which reduced PR responding for fentanyl/sucrose.

**Conclusion:** The three sets of LIFU parameters, with repeated MRgLIFU-NAc sessions, did not produce detectable changes in oral fentanyl/sucrose motivation. Further studies are required to determine effective LIFU parameters needed to reduce NAc-dependent fentanyl motivation.

## Introduction

The opioid crisis remains one of the most significant public health challenges globally [1], with fentanyl and its synthetic analogs driving a steep increase in overdose-related mortality [2]. Chronic opioid exposure induces neuroplastic changes within the mesolimbic reward system, particularly the nucleus accumbens (NAc), which serves as a critical hub for the reinforcing and motivating properties of most misused drugs, including opioids [3,4]. Traditional pharmacological treatments for opioid use disorder (OUD) often suffer from low adherence and high relapse rates [1,5], necessitating the development of targeted, non-invasive therapeutic interventions [6,7].

Transcranial Low-Intensity Focused Ultrasound (LIFU) has emerged as a promising tool for non-invasive neuromodulation [8,9]. Unlike deep brain stimulation that requires surgical implantation, non-invasive LIFU utilizes acoustic energy to modulate neuronal excitability with high spatial precision and deep tissue penetration. LIFU is being tested clinically for efficacy and safety in treating psychiatric disorders, including addiction [10–12]. An open-label pilot study of NAc-targeted focused ultrasound in eight patients with substance use disorders demonstrated the feasibility and potential for long-lasting effectiveness of focused ultrasound to reduce drug craving and drug use [12]. Another clinical pilot study in 10 healthy subjects also demonstrated that 20 minutes of concurrent focused ultrasound/functional MRI (FUS-fMRI) scans safely and transiently altered NAc activity, leading to increased functional connectivity measured with resting-state fMRI [13]. Furthermore, a larger clinical study in 26 healthy adults used FUS targeting the NAc, dorsal anterior cingulate cortex (dACC) or sham, concurrently with fMRI, and showed selective effects on learning and decision making, comparable to that found with deep brain stimulation (DBS) [14]. These studies collectively demonstrate that FUS is a viable approach for non-invasive deep-brain neuromodulation. However, it remains unclear how FUS-NAc might persistently reduce craving and motivation for drugs.

A few preclinical studies investigated FUS-induced effects on drug-motivated behaviors [15–18]. FUS neuromodulation of NAc [16], as well as infralimbic cortex [17], influenced conditioned place preference (CPP) behaviors when tested immediately after sonication. However, to our knowledge, no studies of drug-motivated behaviors demonstrated the attenuating effects of LIFU lasting beyond 1-2 hours without producing cellular damage. As clinical efficacy hinges on long-term behavioral modification, we sought to test motivation for the opioid, fentanyl, at longer intervals from LIFU treatment.

In this study, we aimed to evaluate whether MR-guided LIFU targeting the NAc (MRgLIFU-NAc) would lead to decreased opioid motivation, and to assess the efficacy of different LIFU parameters for decreasing opioid motivation. We modeled opioid motivation in rats using fentanyl/sucrose self-administration and compared it to sucrose-only self-administration using a progressive ratio (PR) schedule of reinforcement that measures motivation for rewards as work requirements increase. As a positive control condition, we also tested whether motivation for fentanyl/sucrose during PR testing is dependent on the NAc using targeted bilateral pharmacological inhibition of NAc with a GABA agonist mixture (muscimol/baclofen). We used our validated model to evaluate the effects of bilateral MRgLIFU-NAc on opioid motivation measured at 4 or 24 hours after LIFU, to ensure minimal or no post-FUS anesthesia effects. To address the significant challenge of precise acoustic energy delivery to the target NAc, we also used Magnetic Resonance Acoustic Radiation Force Imaging (MR-ARFI) [19–23] to ensure accurate targeting and correlate the relative pressures delivered to NAc with behavioral changes across subjects [24]. We hypothesized that MRgLIFU-NAc, in rats that were trained to self-administer oral fentanyl/sucrose will lead to decreased opioid motivation after the effects of anesthesia has dissipated at 4 hours or 24 hours after the FUS.

## Methods

### Animals

All subjects were wildtype Sprague-Dawley rats (Charles River). Rats were between 8-10 weeks of age upon arrival. Operant training began after at least 6 days acclimatization following arrival at the animal facility. In total N=76 rats were used for this study, half of the rats were female and half were male. Rats were housed under 12:12 hour reversed light:dark cycle and procedures were done during the dark cycle. All procedures were performed in accordance with the “Guide for the care and use of laboratory animals” (8th edition, 2011, US National Research Council) and were approved by the University of Maryland School of Medicine Institutional Animal Care and Use Committee (IACUC).

### Food

All training and testing were performed in food-restricted rats. Rats were initially food-restricted to ∼90% of their free feeding weight and fed daily after behavioral training. Rats were allowed to gain weight over the course of the experiment to maintain weights approximately 10% below that of their estimated free feeding weights. Rats were fed standard rat chow.

### Behavioral Experiments

#### Equipment

Behavioral experiments were performed in standard rat operant boxes (Med Associates, Fairfax, VT) with grid flooring and two levers on either side of a liquid reward well connected via tubing to a syringe pump for delivery of either sucrose (5%) or sucrose/fentanyl rewards (5% sucrose / 70ug/ml fentanyl citrate). A red house light on the back wall provided illumination throughout the sessions.

#### Self-Administration Training

Lever presses on the active lever at requisite fixed-ratio (FR) levels produce a 50 µl liquid reward delivered over 1.8 s, together with a 5 s compound cue consisting of a tone stimulus (Sonalert, Med Associates, Fairfax, VT) and white light stimulus illumination above the active lever. Lever responses during this 5 s reward and cue period were recorded but not counted toward completion of the next FR requirement. Inactive lever responses had no programmed consequences and were recorded to determine discrimination between levers. Each training session lasted 150 minutes or until a cap on the maximum number of rewards was met. Reward caps were used in early training sessions to limit potential individual or sex differences during acquisition, as well as potentially adverse effects from overconsumption of fentanyl, and were sequentially set at 15, 30, or 50 rewards per session. Sucrose-alone rats were limited to 500 rewards per session during all FR sessions. Rats were trained for 15-19 FR1 sessions before being moved to higher fixed ratio schedules (FR2 & FR3) for additional training. Rats completed 2-4 baseline progressive ratio (PR) training sessions before LIFU manipulations, which started after a total of 23-26 total FR training sessions. In PR sessions, the response requirement increased after each reward + compound cue delivery. The ascending PR response requirements were 1, 2, 4, 6, 9, 12, 15, 20, 25, 32, 40, 50, 62, 77, 95, 118, 145, 178, 219, 268. Rats that did not consistently earn an average of 10 or more rewards per session at FR3 by the end of training were excluded from the study (n=12 out of 63 sucrose/fentanyl rats and n=1 out of 13 sucrose rats).

#### MR-Guided LIFU Treatment

Following the establishment of baseline drug-seeking behavior, animals were assigned to MRgLIFU-NAc treatment or anesthesia-only control groups (SHAM). Assignment was counterbalanced for sex, average baseline FR3 responses, and average baseline PR responses.

During MRgLIFU-NAc procedures, rats were anesthetized with 1.5%-2.0% vaporized isoflurane and 100% oxygen (flow rate = 1 L /hr) delivered through a nose cone. Breathing rate and body temperature were monitored using an MRI-compatible monitoring system (Small Animal Instruments Inc., Stony Brook, NY). The hair on the scalp was shaved before applying ultrasound gel to ensure adequate acoustic coupling and the transducer was positioned over the animal’s head.

MRgLIFU-NAc was performed using a 7T small animal MRI scanner (BioSpec 70/30, Bruker, Germany) equipped with an MR-compatible 1.5 MHz FUS system (Image Guided Therapy or IGT, Pessac, France). The setup included an 8-element annular array transducer for focused sonication and a single-loop transmit/receive RF coil.

Following positioning of the animal and FUS system at the magnet isocenter, an initial adjustment of the FUS transducer position was made by acquiring axial, coronal, and sagittal T2-weighted spin echo images (TR = 3000 ms, TE = 35 ms, slice thickness = 1 mm, FOV = 40 × 40 mm², matrix = 128 × 128), (**Figs. 1A-C**). The IGT software package was then used to predict and display the location of the FUS focal point relative to the brain, and the mechanical and electronic shifts needed to move the focal point to the left or right NAc were applied (**Fig. 1D**).

**Figure 1:**
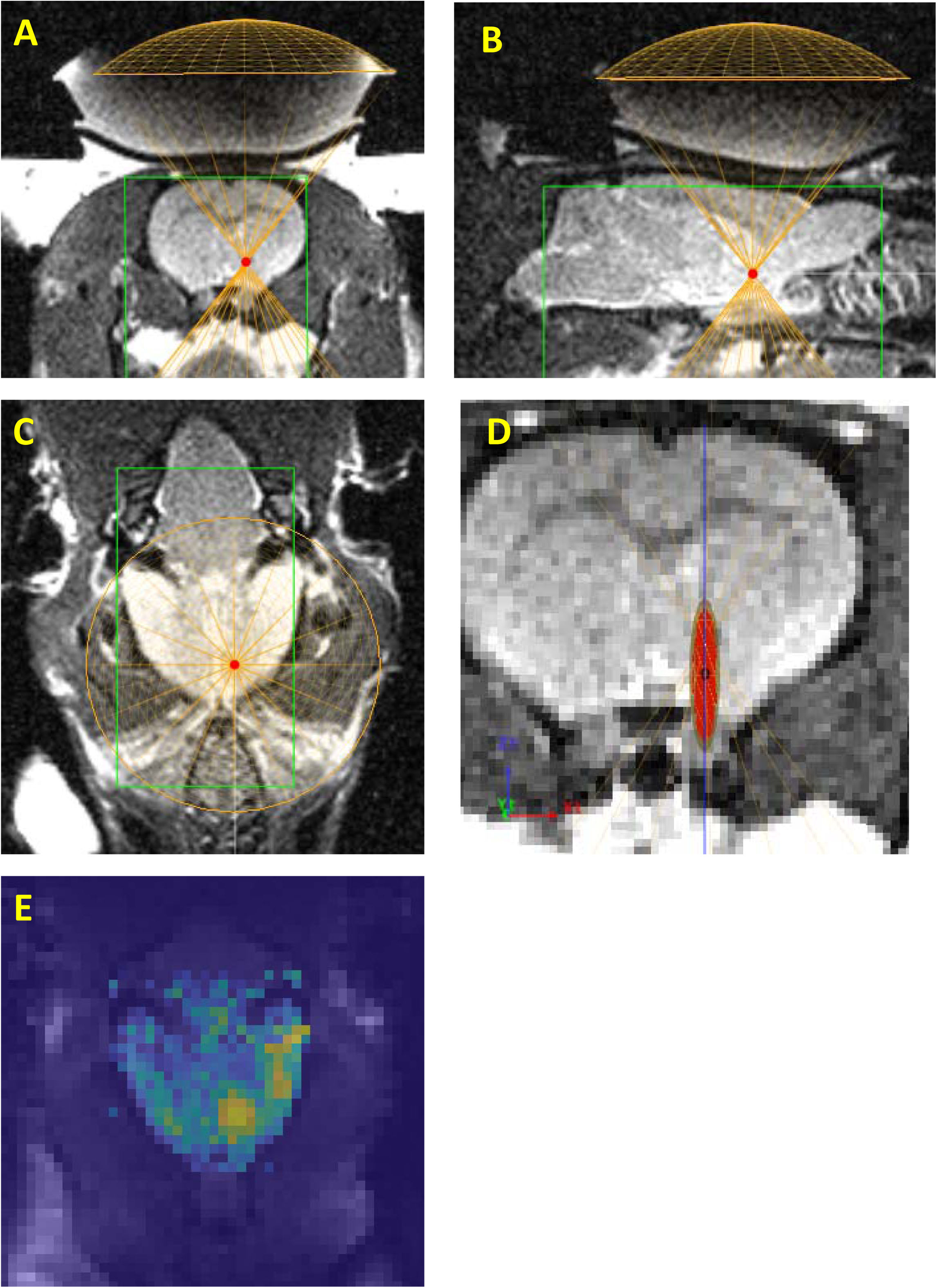
MR-guided targeting and MR-ARFI verification of FUS delivery to NAc. (A–C) Representative axial, coronal, and sagittal T2-weighted MR images used to position the FUS transducer and identify the target anatomy. (D) IGT software visualization of the predicted FUS focal point relative to the brain. (E) Representative MR-ARFI displacement map acquired to verify the location of the FUS focal point.

Next, MR-ARFI was acquired to verify focal targeting (**Fig. 1E**). If the initial focal point was incorrect (i.e., not accurately localized within the NAc), the targeting of the FUS system was refined by adjusting the focal point either electronically along the beam direction or radially by mechanically moving the FUS transducer horizontally above the head. Then, the MR-ARFI acquisition was repeated to ensure that the NAc region was accurately targeted. The MR-ARFI was also used to compare the relative pressure delivered at the foci across different experiments, as explained below. The MR-ARFI acquisition was conducted with the following parameters: TE = 25 ms, TR = 1000 ms, field of view (FOV) = 50 × 50 mm², a matrix size of 64 × 64, and a slice thickness of 2 mm. The duration of each lobe in the bipolar gradients was 4 ms, with a maximum strength of 135 mT/m. The ultrasound pulse duration was fixed at 10 ms to encompass the two central lobes within the bipolar gradient. For all ARFI acquisitions, the input FUS pressure was held constant.

##### Sonication Schemes

Since this study also aims to evaluate the optimal FUS parameters, three sonication schemes were utilized in the LIFU procedures (**Table 1**). In Scheme 1, animals received sonication in a burst mode (500 ms bursts repeated every 5000 ms). Pulses within the bursts were 5 ms in duration with a repetition frequency of 100 Hz. Sonication was applied for 15 minutes at a target acoustic pressure of 2.0 MPa. The spatial peak temporal average intensity (*I_spta_*) in this scheme was 6.3 W/cm^2^.

**Table 1.**
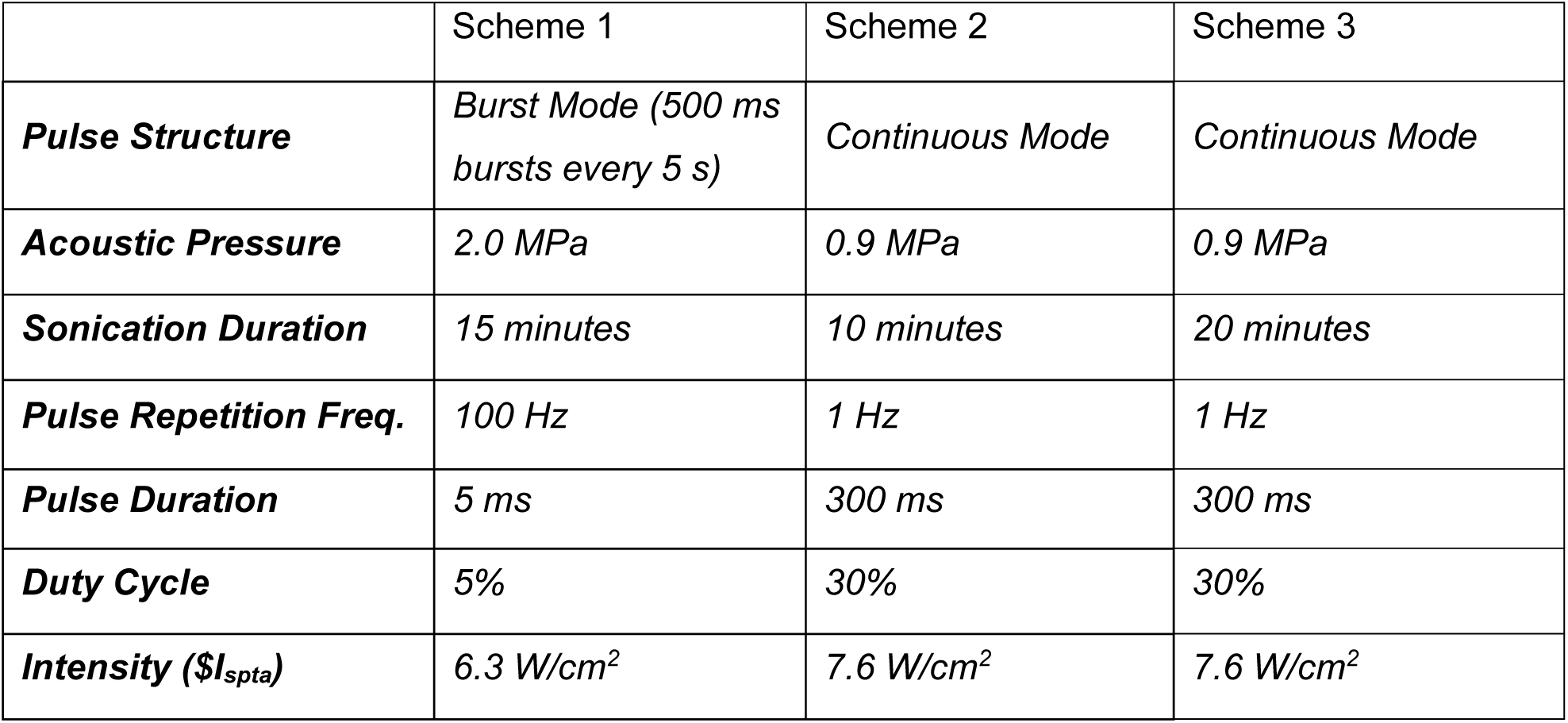
Summary of Three LIFU Schemes for the MRgLIFU-NAc Experiments.

Schemes 2 and 3 both used continuous pulsing but differed in the total duration of sonication. In both schemes, pulses were delivered at a pulse repetition frequency of 1 Hz, with a pulse duration of 300 ms (30% duty cycle) and no secondary burst structure, at a target acoustic pressure of 0.9 MPa (*I_spta_* of was 7.6 W/cm^2^). Scheme 2 consisted of 10 minutes of sonication, corresponding to 600 pulses, whereas Scheme 3 consisted of 20 minutes of sonication, corresponding to 1200 pulses.

#### MR-ARFI image reconstruction and analysis

To verify the location of the focal point and to compare the variation in the displacement due to variation in the delivered FUS pressure across different animal experiments, ARFI displacement maps were generated as follows. First, the brain was segmented by thresholding the magnitude images, resulting in a brain mask. Next, the phase of the ARFI images was unwrapped within the brain mask. The reference phase, obtained from the set of slices with sonication turned off, was then subtracted from the ARFI phase images to eliminate phase variations due to magnetic field inhomogeneities and eddy currents. Finally, the phase difference images were converted to displacements *d* using the formula:

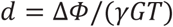

where *ΔΦ* is the MR-ARFI phase difference, *γ* is the proton gyromagnetic ratio, *G* is the amplitude of the encoding gradients, and *T* the total duration of two gradient lobes. The average displacement at the focal point was then calculated in 3 x 3 pixel region (or 2.3 x 2.3 mm^2^) encompassing the peak.

#### Behavioral Testing Post-MRgLIFU-NAc

Rats were tested in PR sessions either at 4 hours or 24 hours after the onset of the MRgLIFU-NAc procedures. Rats were also tested in FR3 sessions at variable delays (indicated in figure timelines) following the post-MRgLIFU-NAc PR test. Rats were fed daily rations immediately after the MRgLIFU-NAc procedures (except for the 4-hour tests) and were therefore similarly food restricted during training and post-MRgLIFU-NAc testing. Individual assignments into MRgLIFU-NAc or Sham groups were consistently maintained across multiple rounds of MRgLIFU-NAc and PR testing for cohorts 1 and 2, whereas a within-subject design was used for cohort 3.

#### Surgical Procedures

Following behavioral training, and post-MRgLIFU-NAc testing in which no effects of LIFU were observed, a subset of rats (N=14) were implanted with bilateral guide cannulas targeting the NAc core to assess whether inactivation of NAc would lead to behavioral changes. Rats were anesthetized with isoflurane, and guide cannulas were targeted to 1 mm above final target to account for injector extending 1 mm beyond cannula tip. Stereotaxic coordinates used to implant guide cannula, relative to skull bregma, were: AP: +1.7 mm; ML: +/− 2.4 mm at a 9 degree angle; DV: 6.1 mm. Cannulas were fixed in place with dental cement and blocked with a dummy cannula until use. Carprofen (5 mg/kg, s.c.) and lidocaine (1 mg/ml; 0.5ml, s.c.) were administered prior to surgery. Rats were given carprofen (5mg/kg, s.c.) for 3 days following surgery. Animals recovered for 1 week following surgery before additional baseline retraining and subsequent PR testing was performed.

#### Histology

Rats that underwent the NAc inactivation study with muscimol/baclofen were transcardially perfused with 0.1M PBS (pH 7.4) followed by 4% paraformaldehyde in PBS. Brains were removed and postfixed in 4% paraformaldehyde overnight and saturated with 30% sucrose before sectioning on a cryostat and imaged at low magnification to determine the location of the cannulas. Rats that had either of the bilateral cannula outside of NAc core were excluded from analysis (n=6). All injection tip locations are shown.

#### Statistics

Comparisons between sucrose and sucrose/fentanyl rats, male and female rats, and groups in the muscimol/baclofen study were made using standard parametric statistical methods (two-tailed t-tests, paired or unpaired as appropriate). Experiments with consistent group sizes across multiple measurements were compared using 2-Way, mixed ANOVAs with repeated measures within the two groups. In cohorts 1 and 2, group sizes varied slightly across timepoints, so mixed linear models were applied with a random intercept for subject. In R, lme4 and lmerTest were used to specify the model and assess overall group-by-time significance, and emmeans were used to obtain estimated marginal means and model-based contrasts at each timepoint. Correlations between MR-ARFI-measured displacements and post-MRg-LIFU-NAc behavioral effects were made using Pearsons linear correlations where Log_2_ Fold Change was calculated from pre-MRg-LIFU-NAc baselines and post-MRg-LIFU-NAc PR data. Significance was taken as p<0.05 for all experiments, and multiple testing was controlled in ANOVAs by controlling the FDR [25].

#### Drugs

Fentanyl Citrate (Cayman Chemical, Ann Arbor, Michigan) was dissolved in water and diluted to final working concentration in sucrose solution. Concentration is given as the weight of the salt. Muscimol (Sigma, Burlington, MA) and baclofen (Tocris, Bristol, UK) were dissolved in sterile saline vehicle and solutions were combined for a final concentration of 100ng/ul for each drug. Muscimol/baclofen mixtures and vehicle were administered under gentle manual restraint at a speed of 0.2 µL per minute at a total volume of 0.5 µL per side resulting in total drug dose of 50 ng baclofen/side and 50 ng muscimol/side. Intracranial cannulae were left in place for an additional 1 min following injection and removed immediately before behavioral testing.

## Results

### Oral Fentanyl/Sucrose Model

Rats were trained to self-administer either oral sucrose (5%) or oral fentanyl citrate mixed with sucrose (70ug/ml fentanyl/5% sucrose) over the course of 21-23 FR training sessions (**Fig.2A**). The pattern of responding for fentanyl/sucrose solutions indicate that rats typically undergo several distinct bouts of consumption and drug ‘satiety’ over the course of the session, similar to that observed during IV fentanyl administration [26] (Figure **2A****, inset**). In fentanyl/sucrose rats, males responded more than females during FR3 sessions (p=0.025, t-test; **Fig. 2B**), but there was no difference in responses between the male and female rats during PR baseline sessions (p=0.89; t-test; **Fig 2C**).

**Figure 2:**
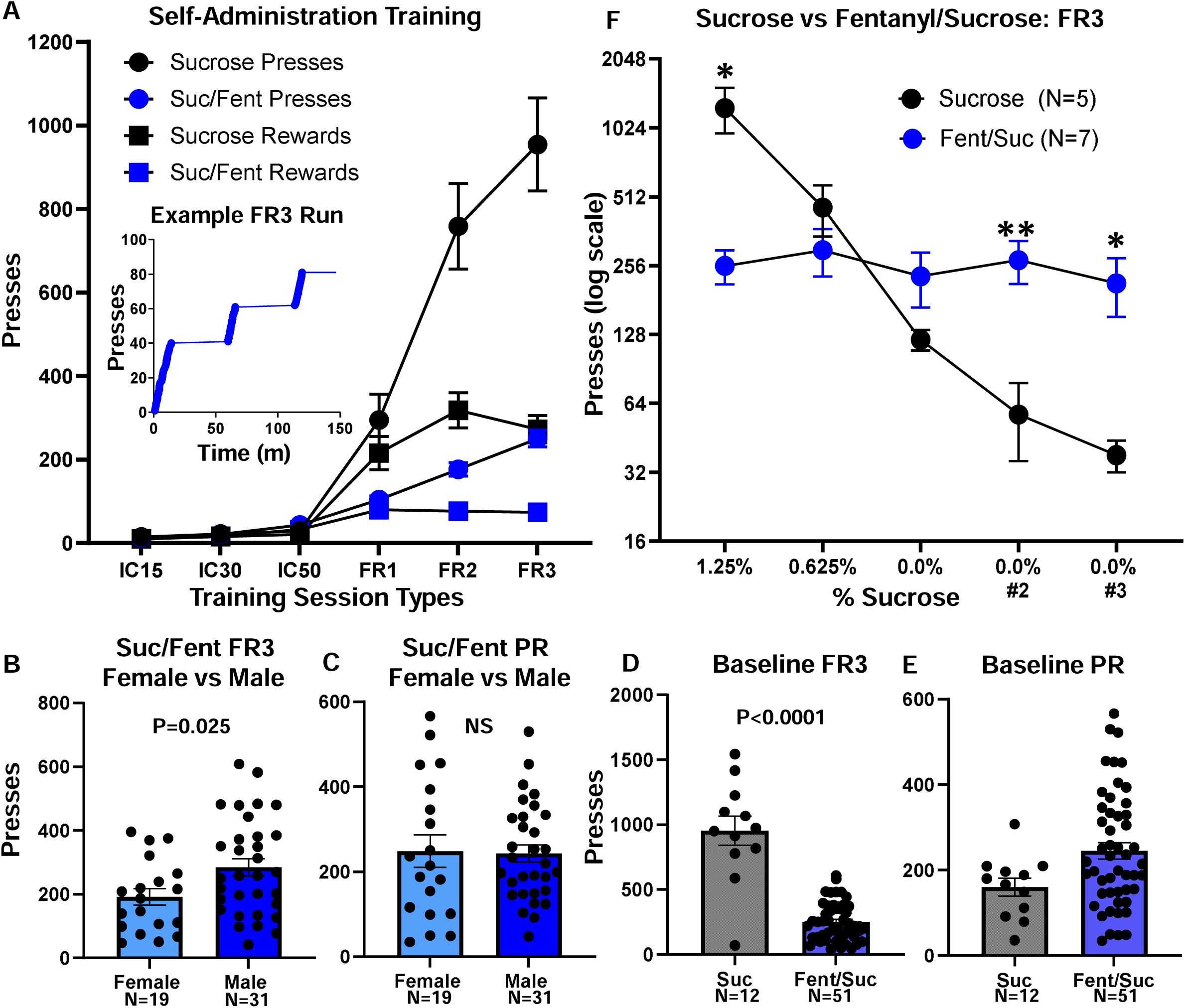
Self-Administration Data. (A) Performance across initial training for fentanyl/sucrose rats (blue) and sucrose alone rats (black). Multiple sessions of each session-type were averaged for each individual. Inset shows the typical pattern of responding during fentanyl/sucrose sessions. (B) Average performance during FR3 training sessions for males and females. (C) Average performance during PR training sessions for males and females. (D) Average performance during FR3 training sessions for sucrose alone vs. fentanyl/sucrose. E) Average performance during PR training sessions for sucrose alone vs. fentanyl/sucrose. P=0.041 for (E), but see text for discussion of this subplot. Statistical results for (B-E) were made using unpaired t-tests. (F) Comparison of sucrose alone vs. fentanyl/sucrose FR3 performance as sucrose concentrations were lowered. Statistics were performed using a 2-Way ANOVA with repeated measures and FDR controlled post-hoc tests. “*” and “**” indicate post-hoc significance to p<0.05 and p<0.01, respectively.

In FR3 sessions, rats responded more for sucrose compared to fentanyl/sucrose solutions (p<0.0001; t-test; **Fig. 2D**). In contrast, during baseline PR testing, rats responded somewhat more for oral fentanyl/sucrose solutions than for sucrose alone (p=0.041; t-test; **Fig. 2E**), however, a subset of sucrose-alone rats consumed the entirety of the sucrose syringe (20 ml) during FR3 sessions, potentially influencing their PR behavior. Therefore, in a subset of sucrose-alone rats (n=5), we used larger (50 ml) syringes throughout FR3 training. We compared the responding of these sucrose-alone rats to highly trained fentanyl/sucrose rats in FR3 sessions where we successively lowered sucrose concentrations from 5% to 0%, remaining at 0% sucrose for 2 additional sessions (**Fig. 2F**).

As expected, all sucrose-alone rats reduced their responding drastically as sucrose concentrations were lowered, however, most fentanyl/sucrose rats maintained a relatively stable level of responding (2-Way ANOVA, Concentration X Group Interaction: p=0.0002), exhibiting significantly greater responding than sucrose-alone rats during the last two 0% sessions (Session 4: p=0.009; Session 5, p=0.028; FDR controlled post-hoc tests following 2-Way Anova) (**Fig. 2F**). Together, these data suggest the fentanyl component of the sweetened drug mixture motivated the PR responding at high costs, but limited the FR3 consumption relative to sucrose-alone at low costs due to drug satiety effects.

### MRgLIFU-NAc at Low-Intensity (MRg-LIFU-NAc) on Fentanyl/Sucrose responding

After baseline FR and PR training, three separate cohorts were treated with either anesthesia-alone or MRg-LIFU-NAc bilaterally. **Figs. 1A-1D** demonstrates the positioning of the transducer relative to the brain using IGT software. **Fig. 1E** is an example ARFI displacement map demonstrating the spatial distribution of calculated displacement values. Two cohorts (#1 & #2; **Figs. 3 & 4**) underwent multiple rounds of MRg-LIFU-NAc followed by PR testing 24 hours post-MRg-LIFU-NAc. Additional FR3 training was also conducted between MRg-LIFU-NAc treatments, as indicated in **Figs. 3A & 4A**. Detailed sonication parameters used for each of the MRg-LIFU-NAc rounds are shown in **Table 1**.

**Figure 3:**
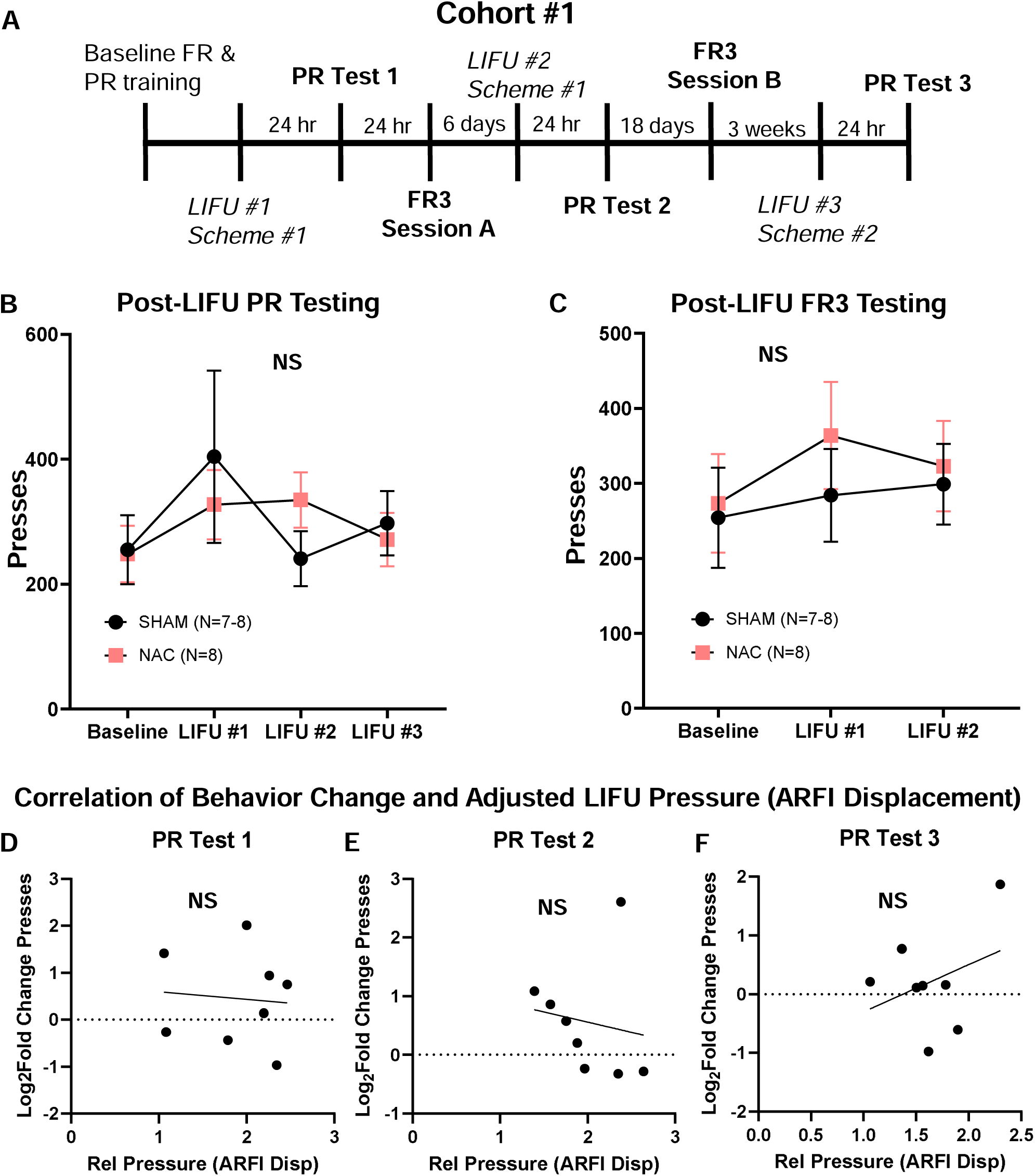
Effects of Focused Ultrasound on Fentanyl/Sucrose behavior in Cohort #1. (A) Timeline of Experiment. (B) Baseline and post-focused ultrasound PR results. (C) Baseline and Post-focused ultrasound FR3 results. Statistics for (B-C) used mixed linear models where “NS” Indicates no significant interaction was found between treatment and session overall (Ps>0.34). Pairwise contrasts between treatment groups were also performed at each post-focused ultrasound session, and all p’s>0.39 for (B-C). (D-F) Correlations between subject-level ARFI displacements and behavioral changes relative to baseline averages. Statistics used Pearson’s correlations.

**Figure 4:**
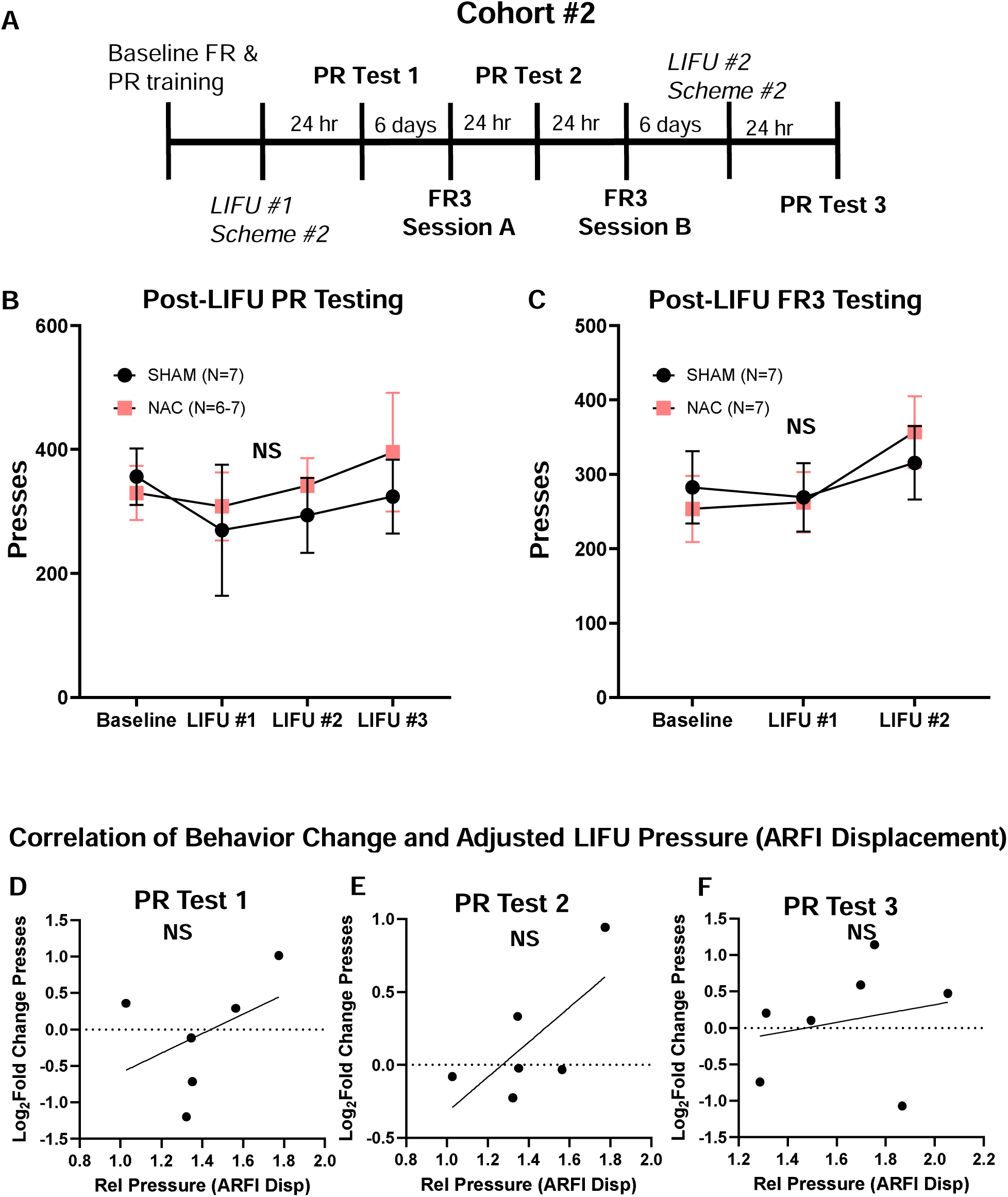
Effects of Focused Ultrasound on Fentanyl/Sucrose behavior in Cohort #2. (A) Timeline of Experiment. (B) Baseline and post-focused ultrasound PR results. Statistics for (B) used mixed linear models where “NS” Indicates no significant interaction was found between treatment and session overall (TreatmentXSession F(3,35.31)=0.31; p=0.81). Pairwise contrasts between treatment groups were also performed at each post-focused ultrasound session, and all p’s>0.45 for (B). (C) Baseline and Post-focused ultrasound FR3 results. “NS” indicates no overall interaction between Treatment and Session (TreatmentXSession F(2,24)=2.67; p=0.09; Two-Way ANOVA). Post-hoc tests at each timepoint reveal no significant differences at any time between groups (all p’s>0.52). (D-F) Correlations between subject-level ARFI displacements and behavioral changes relative to baseline averages. Statistics used Pearson’s correlations.

The training and sonication timeline for Cohort 1 is shown in **Fig. 3A**. In this cohort, after the 1^st^ round of MRg-LIFU-NAc using Scheme #1, we detected no significant group differences between MRg-LIFU-NAc and controls in the post-MRg-LIFU-NAc PR test sessions (**Fig. 3B**). We also did not observe any changes in behavior during an FR3 session conducted 48 hours after MRg-LIFU-NAc (**Fig. 3C**). We performed a 2^nd^ round of MRg-LIFU-NAc using Scheme #1 parameters, 1 week after the 1^st^ round, and again saw no differences in PR responding between the rats that had MRg-LIFU-NAc and control rats 24 hours later (**Fig. 3B**), nor during a follow up FR3 session (**Fig. 3C**). 3 weeks later, we performed a final 3^rd^ round of MRg-LIFU-NAc in this cohort using Scheme #2 parameters, and detected no change in PR responding between rats that received MRg-LIFU-NAc and control rats that had the same duration of anesthesia (**Fig. 3B**). We further analyzed these data at an individual level to correlate behavioral changes during post-MRg-LIFU-NAc PR testing with MR-ARFI-measured displacements achieved during FUS. We observed no significant relationships with MR-ARFI displacement and changes in behavior relative to baseline (**Figs. 3D, E & F**).

The training and sonication timeline for Cohort 2 is shown in **Fig. 4A**. In Cohort 2, we again tested the effects of MRg-LIFU-NAc against anesthesia-only controls on their performance 24 hours later. We observed no difference in PR responding at 24 hours after the first MRg-LIFU-NAc session using Scheme #2 (**Fig. 4B**), nor during a follow up PR test one week after this first session (**Fig. 4B**). An FR3 session between these two PR tests after the first MRg-LIFU-NAc session also showed no difference in performance between controls and LIFU-treated rats (**Fig. 4C**). We performed a second session of MRg-LIFU-NAc in this cohort using Scheme #2, and again, we observed no changes in PR performance 24 hours later (**Fig. 4B**). Correlational analyses between ARFI-measured tissue displacement and performance change relative to baseline revealed no statistically significant trends in any tests during this 2^nd^ cohort (**Figs. 4D, E, F**).

Cohort 3 used a continuous sonication scheme (Scheme #3), using a within subject design, and shortened 4-hour interval between MRg-LIFU-NAc and PR testing. Half the rats received MRg-LIFU-NAc the first week and the other half received Sham MRg-LIFU (anesthesia only) the first week; the animals were then switched into the opposite condition approximately 1 week later (N=14). We again observed no changes in PR performance at 4 hours, nor in FR3 performance at 24 hours post-MRg-LIFU-NAc (**Figs. 5B,C**).

**Figure 5:**
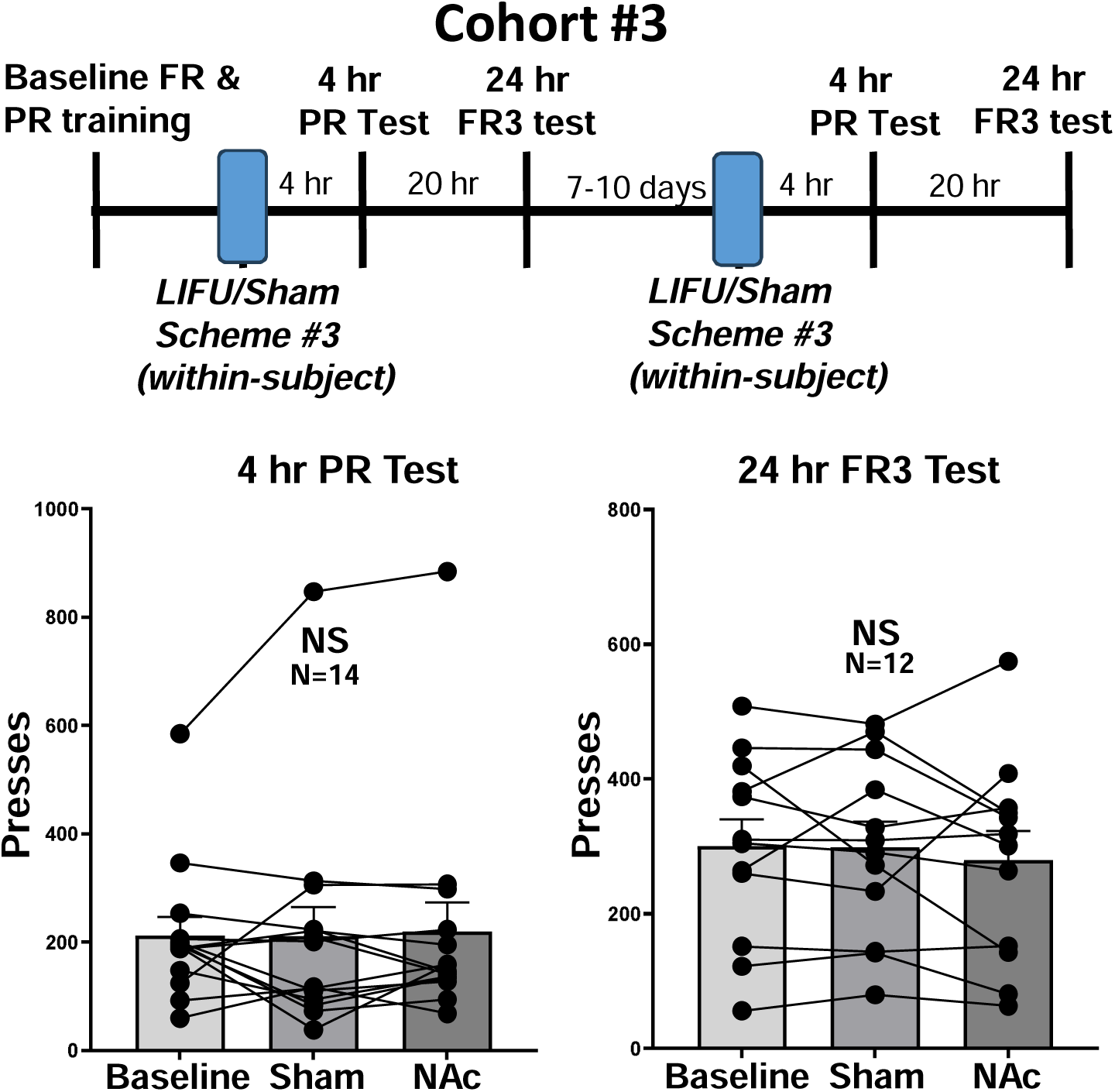
Effects of Focused Ultrasound on Fentanyl/Sucrose behavior in Cohort #3. (A) Timeline of Experiment. (B) Baseline and post-focused ultrasound PR results. “NS” indicates no significant group effect: F(2,18)=0.375; p=0.69, 1-Way ANOVA w/ Repeated Measures. (C) Baseline and post-focused ultrasound FR3 results “NS” indicates no significant group effect: F(2,14)=0.147; p=0.864, 1-Way ANOVA w/ Repeated Measures.

### Oral Fentanyl/Sucrose PR responding is NAc dependent

A subset of rats was implanted with bilateral infusion cannulae targeting the NAc (**Fig. 6A**) and tested in PR for fentanyl/sucrose responding. In a counterbalanced, within-subject design, pharmacological inactivation with muscimol/baclofen (50ng+50ng/0.5ul/side) resulted in a significant reduction of PR responding (p=0.004, paired t-test, **Fig. 6B**) relative to saline infusions. These data indicate that PR responding for sweetened oral fentanyl is NAc-dependent.

**Figure 6:**
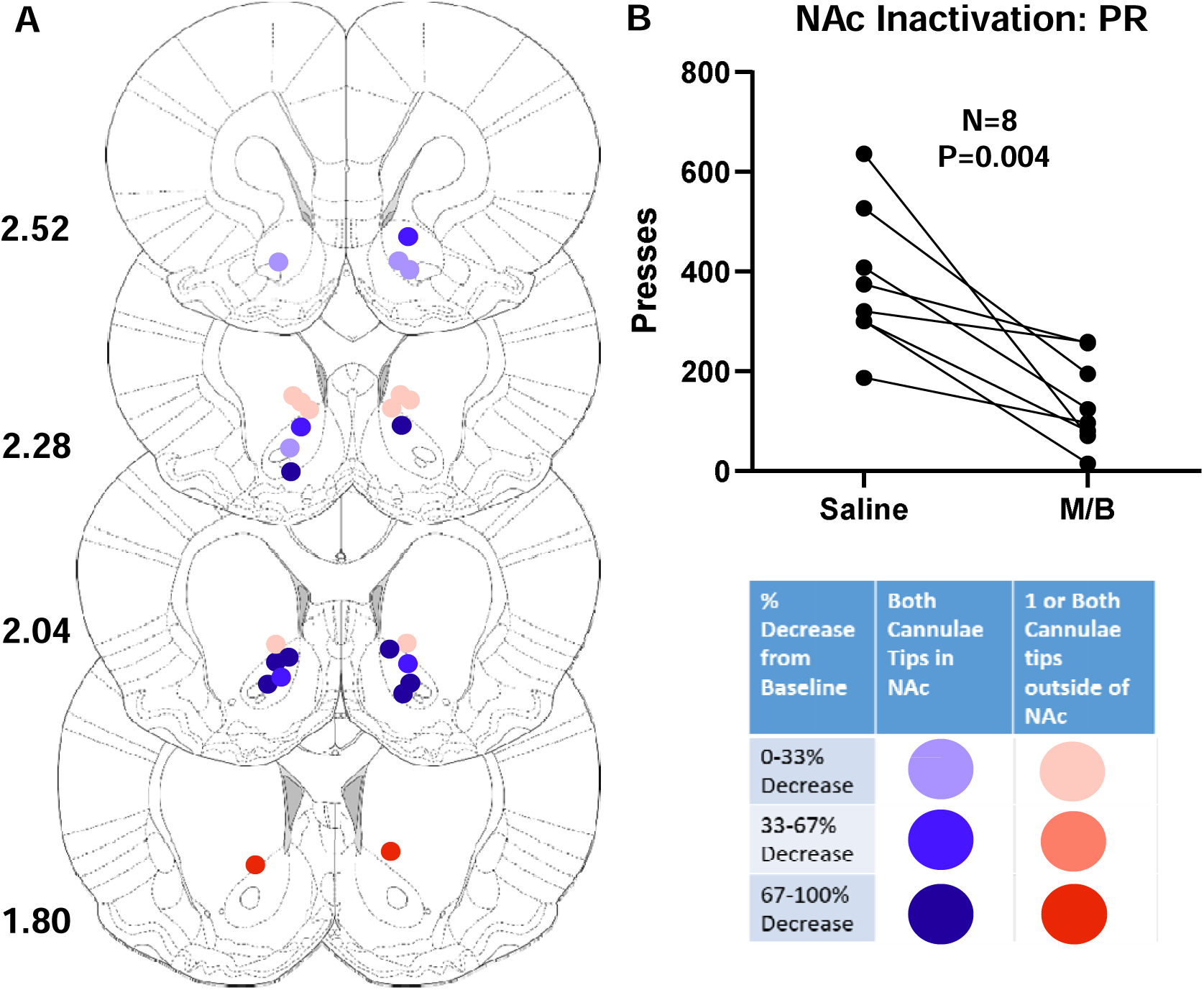
Intracranial Muscimol/Baclofen Reduces PR Responding. (A) Locations of injector tips colored according to % decrease from saline responding. Only placements with both injector tips in NAc were included in statistical analysis. (B) PR responding for saline vs muscimol/baclofen (50ng/ul for both compounds; 0.5 ul/side). Statistical comparison used a paired t-test.

## Discussion

This study aimed to evaluate whether MRg-LIFU-NAc produces sustained changes in motivation for oral fentanyl/sucrose self-administration behavior. We performed the following: 1) We optimized and used an oral fentanyl self-administration and PR model that we validated as sensitive to pharmacological inhibition of NAc. 2) We performed the MRg-LIFU-NAc in three cohorts of rats that were trained to self-administer oral fentanyl/sucrose, and tested three different LIFU schemes to test effects from different FUS parameters. 3) We tested whether repeated MRg-LIFU-NAc, up to 3 sessions, in some of these rats have cumulative effects on opioid motivation; 4) We assessed the animals at different time points after the MRg-LIFU-NAc sessions, at 4 hours and again at 24 hours later, to minimize any potential confounds on the opioid motivation from post-anesthesia effects. 5) We used an MR-ARFI technique to ensure accurate targeting of the bilateral NAc and evaluate the relative focused ultrasound energy delivered to the NAc (ARFI displacement). Despite these multiple testing and trials using different parameters, we found that motivation for sweetened oral fentanyl was not significantly altered at 4 or 24 hours after MRg-LIFU-NAc under any of the conditions we tested. We also found no behavioral changes in FR3 behavior at longer intervals after MRg-LIFU-NAc.

### Oral Opioid Motivation Model

Preclinical models of opioid motivation using oral drug delivery have practical advantages over IV models, such as not requiring catheterization surgeries nor catheter patency maintenance. Although delayed peak drug levels and potentially aversive taste qualities are challenges to oral delivery, oral oxycodone and oral fentanyl models have been used previously in rats and mice [27–30]. In early pilot experiments after limited fentanyl/sucrose training, we found that removal of sucrose from fentanyl/sucrose solutions resulted in substantial reductions in fentanyl consumption (data not shown). However, after extensive training (>30 total sessions), the majority of sucrose/fentanyl rats we tested maintained relatively stable levels of fentanyl consumption after sucrose was tapered to 0%. This suggests that the robust level of lever pressing in fentanyl/sucrose rats during PR tests (where the price for successive rewards rapidly escalates), is motivated in part by opioid reinforcement.

We note that rats do not reach fentanyl satiety in a PR session, as they earn fewer reinforcers in a PR session than they earn in a single consumption bout during FR3 sessions. We demonstrate that high motivation for fentanyl/sucrose requires NAc activity, as pharmacological inhibition of the NAc with a GABA agonist mixture (muscimol/baclofen), significantly blunted motivation for fentanyl/sucrose. Therefore, the oral sweetened fentanyl PR test in well trained rats is an appropriate model to test for potential effects of MRg-LIFU-NAc to inhibit opioid motivation, with the caveat that caloric motivation likely also contributed to task performance in this model (since rats were food restricted prior to testing).

### MR-ARFI Targeting

MRgLIFU targeting the NAc in a rodent is technically challenging due to the small size and shape of the structure, as well as variations in skull thickness between sexes and individuals. Sonic aberration by the skull can lead to reduced accuracy and inconsistency in delivering energy to the foci of the transducer system [31,32]. We previously demonstrated that MR-ARFI displacement is a strong predictor of FUS power delivered to target, and that re-positioning of the transducer based on MR-ARFI generated displacement maps leads to more accurate targeting of acoustic energy [24]. Nevertheless, significant variation in displacement at the NAc persists, as we measured a range of relative pressures varying by about two-fold across subjects. However, we did not find significant correlations between the FUS pressure delivered to NAc and differences in opioid motivation relative to baseline.

### Previous Studies of Focused Ultrasound on Drug-motivated Behavior

To our knowledge, the work shown here is the first study of focused ultrasound effects on operant drug self-administration behaviors. Three previous studies measured LIFU effects on experimenter administered drug conditioned place preference (CPP) behavior at short intervals following sonication [15–17]. The first of these studies established a degree of morphine CPP over several sessions of training before splitting rats into sham and sonication groups, and administered LIFU treatment prior to 10 additional CPP training sessions [15]. Although LIFU did not reduce CPP, the NAc sonication group did not develop further place preference for the morphine paired side over the additional training sessions, whereas the sham group did [15]. Since the training sessions took place immediately following LIFU, any effects on CPP learning might have been short lived [15].

Another study demonstrated that infralimbic cortex stimulation with LIFU immediately prior to testing reduced methamphetamine-primed CPP reinstatement after CPP extinction sessions [17]. However, the authors found no changes in CPP when the rats were assessed a week later [17], indicating that the effects from LIFU were not long-lasting.

More recently, LIFU targeting the NAc was shown to reduce fentanyl-motivated CPP behavior in rats [16]. Like the other CPP studies, this study differed from ours in several respects, including sonication parameters, transducer design, targeting method, and the behavior tested [16]. Perhaps most importantly, it used a crossover design with individuals undergoing only a brief 2 hour washout between LIFU and sham manipulations, with behavioral testing starting within an hour of completing LIFU. In this study, effects to inhibit fentanyl CPP after LIFU targeting the NAc-were apparently short-lived, since half the subjects also received sham interventions and testing shortly after post-LIFU testing on the same day [16]. Another study in mice showed that heroin CPP was inhibited 24 hours after sonication to the NAc combined with microbubbles [18]. While this study demonstrated longer lasting effects of focused ultrasound in the ultrasound+microbubble group, the ultrasound alone group did not show an attenuation of the CPP behavior [18]. Furthermore, sonication with microbubbles using their protocol led to extravasated erythrocytes and damage to the synaptic ultrastructure measured with electron microscopy, suggesting that NAc damage may be responsible for the extended effects [18].

### Detailed Discussion of Interval between LIFU and Behavioral Testing

Our work tested the effects of LIFU on opioid motivation at 4 or 24 hours post-MRg-LIFU-NAc, as well as later timepoints. It is possible that NAc-targeted LIFU using our parameters does inhibit drug responding, but this inhibition is too short-lived to measure using the methods we employed. In pilot work, we found that our anesthesia-only controls exhibited variable operant behavior for up to 2 hours following anesthesia, precluding operant behavioral testing at very short intervals. It is also plausible that the depth and duration of anesthesia we employ for our protocol might have interacted with LIFU modulation of neural activity. As both LIFU and anesthesia disrupt ion channel function [33,34], they might have interacted in yet-undescribed ways.

Several other human and rodent studies of LIFU-mediated effects demonstrated persistence of effects in the 0.5 hr – 2 hr range beyond the sonication itself [14, 35–37]. For example, LIFU modulation of the prefrontal cortex reduced dopamine release in the NAc [37]. In that study, a two-minute sonication scheme at 13W/cm2 resulted in a reduction in dopamine release in the NAc for 2 hours, and the effect was sonication-intensity dependent, in which higher and lower intensities decreased the magnitude of FUS effects on dopamine [37]. While measured neuromodulatory effects of LIFU were shown to persist beyond sonication and may explain previously observed CPP effects shortly after sonication, LIFU neuromodulatory effects described thus far did not persist long enough to explain the apparently robust and long lasting effects observed in a clinical study [12].

### Sonication Parameters

In this study, we employed three sonication schemes to investigate their impact on fentanyl motivation: a Burst Mode (Scheme 1), utilizing 500 ms bursts of sub-pulses (50% duty cycle, 100 Hz PRF) delivered every 5 seconds, a Continuous Mode (Scheme 2), consisting of 300 ms pulses at a 1 Hz repetition frequency for 10 min., and a similar but longer Continuous Mode using 20 min. sonication duration (Scheme 3). Similar parameters were explored in the literature with varying degrees of success. For instance, Deveci et al. used a burst scheme with a 50% duty cycle but employed a significantly higher PRF of 250 kHz and a fundamental frequency of 2.4 MHz, with bursts lasting 300 ms every 2 seconds. Conversely, Yoo et al. applied 0.5 ms pulses, delivered continuously at a 100 Hz PRF for 30 minutes [16]. While these parameters were similar to our Continuous Mode, a major point of divergence remains the fundamental frequency; our system operated at 1.5 MHz, which is between the 0.6 MHz and 2.4 MHz frequencies used in these two previous studies.

Identifying the optimal LIFU parameters remains a significant challenge, as different neuronal populations respond differently to acoustic energy and in a non-linear fashion [38,39]. Nevertheless, similar acoustic parameters to those used in this work were previously shown to modulate various neuronal types in a dose-dependent manner [40]. The relative lack of effect on fentanyl-seeking behavior in the current study may be attributed to some combination of suboptimal sonication parameters and the delayed testing intervals (4-24 hours), during which any putative transient neuromodulatory effects may have faded. Future research should investigate the interaction between sonication parameters and duration of behavioral effects.

### Conclusion and Future Directions

In summary, across three cohorts of rats, and three different sonication schemes, MRg-LIFU-NAc did not attenuate fentanyl/sucrose in this study. Although clinical pilot studies demonstrated that LIFU might be a safe and effective treatment to reduce craving and ultimately relapse in people with SUD [11,12], recent serious adverse events raised critical safety concerns [41,42]. Therefore, preclinical rodent studies should be performed first to systematically determine the targets and sonication parameters that are required to alter drug-motivated behavior safely. Future work should establish FUS procedures that give rise to lasting behavioral changes, which can then be used to determine mechanisms associated with these changes, before the therapeutic potential of focused ultrasound for OUD and other SUDs can be realized.

